# CLARA: Compound Library Active Learning with Reasoning Agents

**DOI:** 10.64898/2026.10.05.756921

**Authors:** Satya Pratik Srivastava, Vaidehi Saxena, Disha Singh, Harshit Singh, Rajeev Kumar Singh

## Abstract

Batch active learning for ligand discovery requires a recurring decision about whether to exploit high predicted affinity, reduce model uncertainty, or broaden chemical coverage. We test whether a language model can make this strategy-level decision while candidate ranking and label revelation remain controlled by a deterministic Gaussian-process environment. Across four protein targets, ten paired seeds, and a 360-compound assay budget, a GPT-5.6 Sol agent outperformed random sampling in all 40 matched campaigns. Its final top-2% recall was 0.412 on D2R, 0.374 on Mpro, 0.448 on TYK2, and 0.683 on USP7. It was statistically tied with the strongest fixed policy on Mpro and USP7 but underperformed target-specific leaders on D2R and TYK2. The agent shifted from exploration-oriented early batches toward exploitation and produced 686 testable threshold claims with an 86.3% atomic success rate, although 19.8% of forecasts were not numerically scorable. In 200 prespecified information-ablation campaigns, removing uncertainty, diversity, or memory caused only small endpoint changes, while shuffling explanatory evidence reproduced the full allocation path in 39 of 40 paired campaigns. A deterministic 80% availability constraint reduced recall on every target but preserved the target-dependent pattern of wins and losses. These results support language-model strategy control as a feasible and auditable layer.

## 1 Introduction

Experimental ligand discovery is a search problem under a strict measurement budget. Active learning uses observations collected so far to choose the next compounds, ideally concentrating assays on high-value regions without losing the information needed to improve the model. Early work demonstrated that active selection can outperform random screening in drug discovery [6], while later work showed that batch selection introduces a cost of parallelization and makes diversity important [1]. Gaussian-process (GP) surrogates are attractive because they provide both predictions and uncertainty, enabling acquisition rules that trade off exploitation and information gathering [4].

No single acquisition strategy is expected to dominate every chemical landscape. The four-target benchmark that supplies our data and software found substantial dependence on the target, representation, kernel, and acquisition policy [5]. This suggests a meta-control problem: instead of freezing one acquisition mixture for an entire campaign, can an agent alter the mixture after observing campaign progress?

We evaluate a constrained language-model agent that never sees hidden affinities and never directly selects unrestricted molecules. At each cycle it allocates a batch among three deterministic queues: high predicted mean, high posterior uncertainty, and chemical diversity. The design separates language-model judgment from numerical modeling and assay simulation. Our contributions are:

- a paired four-target benchmark comparing 40 agent campaigns with 360 fixed-policy campaigns;
- an auditable protocol that records every allocation, rationale, expected outcome, selected index, and revealed result;
- outcome scoring for testable rationale claims and 200 prespecified information-ablation campaigns; and
- an operational extension in which only a deterministic 80% of post-initial candidates are available.

Figure 1 summarizes the complete experimental loop and the four prespecified analysis tracks derived from its decision logs.

**Figure 1:**
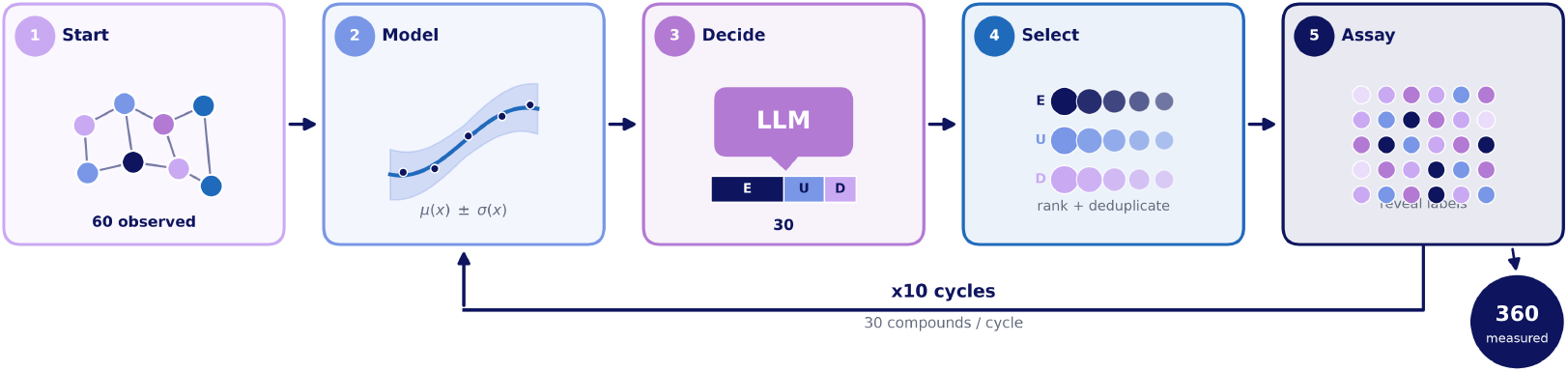
End-to-end experimental workflow. All methods begin from matched initial observations. A GP summarizes the revealed data; the language-model agent allocates each batch across exploitation, uncertainty, and diversity queues; deterministic selection enforces uniqueness and access constraints; and the virtual assay reveals only the selected labels. Ten cycles produce a 360-compound endpoint. The complete decision log supports the core benchmark, next-batch forecast scoring, information ablations, and the operational availability experiment.

## 2 Related work

Active search prioritizes finding members of a valuable class rather than only minimizing predictive error. Drug discovery was an early motivating application [6]. Nonmyopic batch active search formalized the simultaneous selection setting and demonstrated the importance of diversity in batch construction [1]. Bayesian optimization and GP upper-confidence methods provide a complementary view in which posterior mean and uncertainty guide expensive evaluations [4].

Molecular fingerprints encode local structural environments in fixed-length vectors; extended-connectivity fingerprints remain a widely used representation [3]. Similarity kernels, including Tanimoto-style kernels, allow kernel machines and GPs to operate on fingerprint representations [2]. The new element is a language model acting only as the allocation agent, not as the molecular predictor or assay oracle.

## 3 Methods

### 3.1 Datasets and representation

We use four ligand-affinity datasets: D2R (2,502 compounds), Mpro (2,749), TYK2 (9,997), and USP7 (1,799). Each table contains unique, nonempty SMILES and a finite affinity value. Molecules are represented by 4,096-bit Morgan fingerprints with radius 4, consistent with extended-connectivity representations [3]. A small exact GP uses a fixed Tanimoto kernel, observation noise 0.05, and numerical jitter 10^*−*6^.

### 3.2 Campaign protocol

Each campaign begins with 60 randomly measured compounds. Ten cycles add 30 compounds each, producing a final budget of 360. For every target and seed, all methods share the same initial set. After fitting the GP to revealed labels, the environment constructs three ordered queues from the remaining pool:

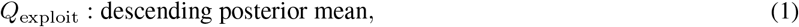

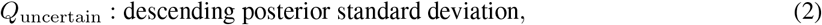

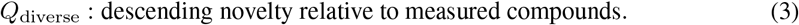

The agent emits nonnegative integers (*n*_*e*_, *n*_*u*_, *n*_*d*_) satisfying *n*_*e*_ + *n*_*u*_ + *n*_*d*_ = 30. The environment then selects unique candidates from the queues with deterministic de-duplication, reveals only their labels, and advances the campaign. Hidden pool labels are structurally absent from the agent state.

### 3.3 Agent information and outputs

The agent receives target and cycle identifiers, the remaining budget, observed performance, GP mean and uncertainty summaries, chemical-space coverage, revealed prior decisions and outcomes, and observed-only fingerprint-enrichment evidence. It returns the three allocations, a rationale, and an expected outcome for the next batch. The core experiment used GPT-5.6 Sol through versioned Codex tasks.

### 3.4 Baselines and evaluation

Nine fixed policies span random, pure and mixed exploit/uncertain/diverse allocations. The primary endpoint is top-2% recall after 360 measurements. We also report top-1% and top-5% recall, simple regret, and the number of unique scaffolds among recovered top-5% compounds. Paired differences use the common target-seed initialization; 95% intervals are percentile bootstrap intervals over the ten paired seed differences.

### 3.5 Forecast scoring, ablations, and availability

Each expected outcome is scored against the immediately following batch. A rule-based parser extracts explicit numerical thresholds into atomic claims. We report the fraction of forecasts that are scorable, atomic-claim success, the mean fraction of clauses satisfied, and conjunction success. Because the forecasts do not contain probabilities, this is not probabilistic calibration.

Five paired information conditions are evaluated over four targets and ten seeds: full information, no uncertainty, no diversity, no memory, and shuffled evidence. Shuffling preserves evidence scores, signs, ranks, and volume while permuting fingerprint-bit identities. The operational benchmark deterministically retains 80% of the post-initial library using canonical isomeric SMILES and a salted SHA-256 mapping independent of affinity. The shared initial 60 are grandfathered to preserve pairing.

## 4 Results

### 4.1 The agent consistently beats random but not every target-specific fixed policy

The agent beat random sampling in all 40 matched campaigns. Mean endpoint top-2% recall was 0.412 on D2R, 0.374 on Mpro, 0.448 on TYK2, and 0.683 on USP7 (Figure 2). The paired improvements over random were 0.266, 0.254, 0.412, and 0.503, respectively; every 95% interval lay above zero.

**Figure 2:**
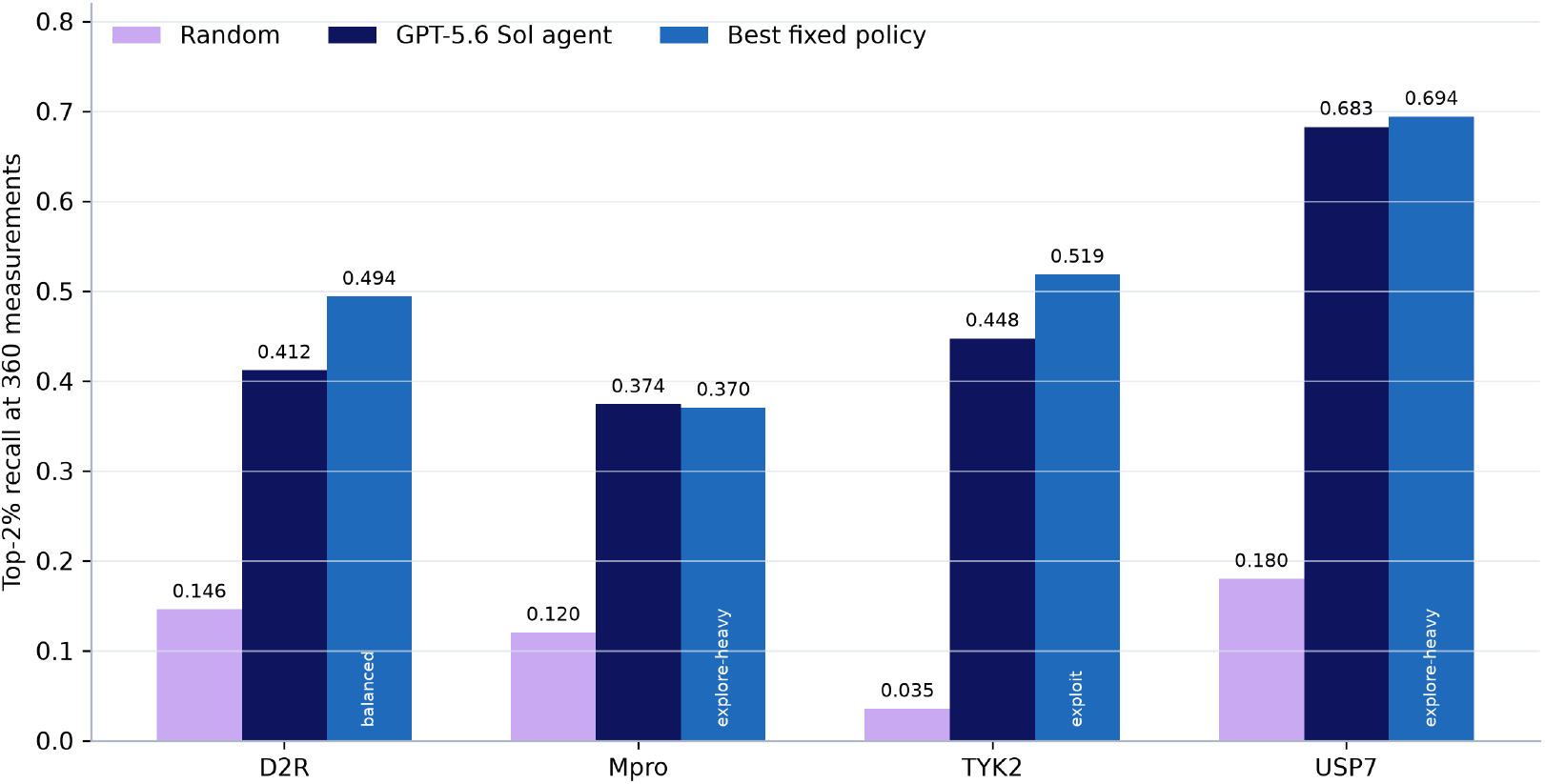
Endpoint top-2% recall after 360 measurements. Bars show means over ten paired seeds. The strongest fixed policy is selected separately for each target. The agent is uniformly better than random but not uniformly better than the target-specific leader.

Against the strongest fixed policy, the result was heterogeneous (Table 1). D2R favored the balanced policy by 0.082 and TYK2 favored pure exploitation by 0.071; both paired intervals excluded zero. Mpro differed from explore-heavy by only +0.004 and USP7 differed from explore-heavy by -0.011, with both intervals crossing zero. The agent found the global-best compound in all Mpro, TYK2, and USP7 runs and seven of ten D2R runs.

**Table 1:**
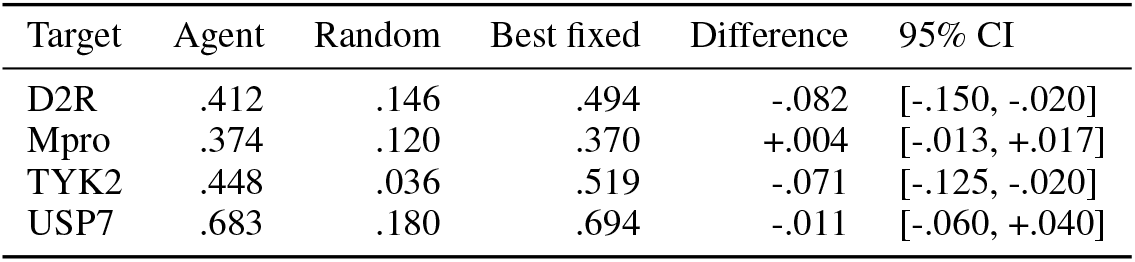
Core endpoint comparison. Values are mean top-2% recall; intervals describe the paired agent-minus-strongest-fixed difference.

Learning curves show that the agent separated from random early and maintained that lead (Figure 3). D2R and TYK2 developed persistent gaps relative to their target-specific fixed leaders, whereas Mpro and USP7 remained close for much of the budget. Endpoint recall therefore does not reflect a single anomalous final batch.

**Figure 3:**
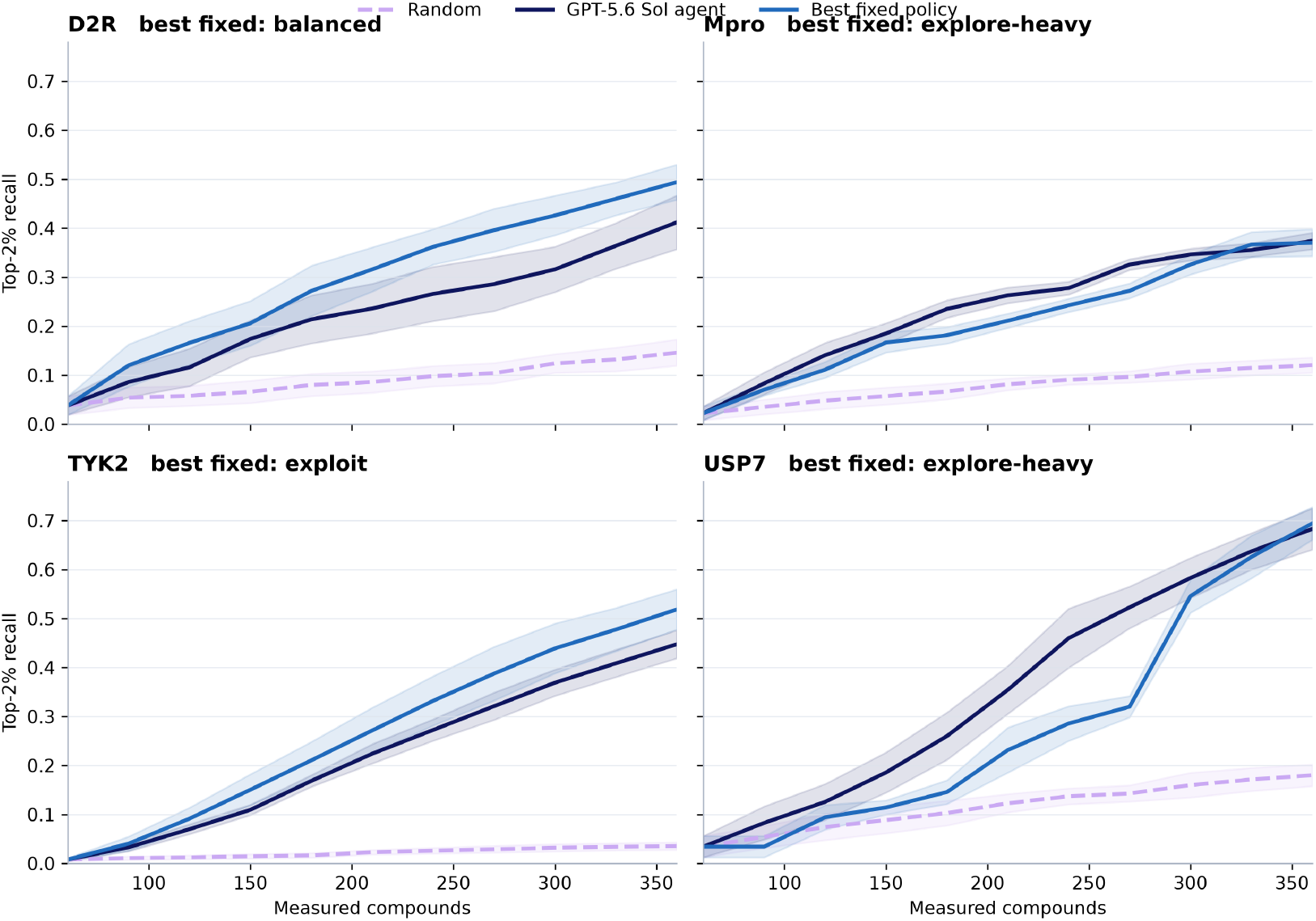
Top-2% recall over the assay budget. Lines are means over ten seeds and bands are approximate 95% confidence intervals based on the standard error.

### 4.2 Allocations adapt across targets and campaign stages

The mean exploit allocation rose from 8.2 compounds in cycle 1 to 18.6 in cycle 10, while uncertainty fell from 11.6 to 6.7 and diversity from 10.2 to 4.7. Target averages reveal distinct regimes: TYK2 was strongly exploitative (19.05 of 30 compounds per batch), Mpro emphasized uncertainty and diversity (5.80 exploit, 12.90 uncertain, 11.30 diverse), and D2R and USP7 used intermediate mixtures. Thus, the agent changed actions with both target and campaign stage. This behavioral adaptation does not establish that the natural-language rationale is a faithful causal explanation.

The comparison with all fixed policies further illustrates this heterogeneity (Figure 4). The agent comfortably beat random, uncertainty-only, and several mismatched mixtures, but target-matched fixed policies remained difficult to beat. The heat map uses the blue family employed elsewhere in the benchmark figures; signed cell labels retain which method is better.

**Figure 4:**
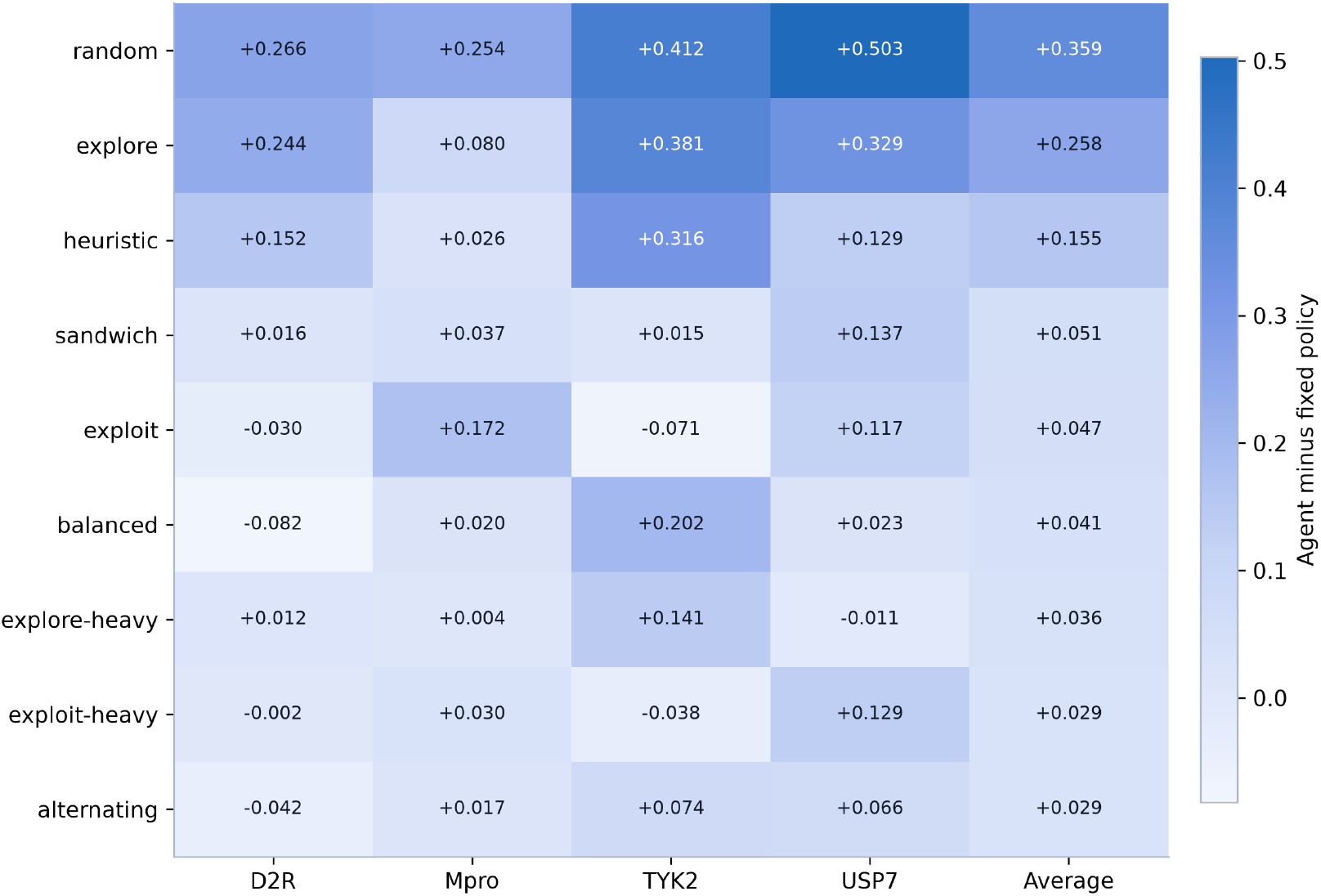
Agent minus fixed-policy mean top-2% recall. Positive values favor the agent. The average column is unweighted across targets and should be interpreted together with target-specific cells.

### 4.3 Expected outcomes are often correct when testable

Among the 400 expected outcomes, 321 contained at least one numerically supported threshold, while 79 (19.75%) were unscorable. The parser identified 686 atomic claims, of which 86.30% were successfully verified. On average, 87.28% of the clauses within each decision were satisfied, and all clauses were satisfied simultaneously in 71.65% of scorable decisions. Scorability was 100% for D2R and TYK2, 83% for Mpro, and 38% for USP7, where a substantial fraction of forecasts were qualitative. These results indicate that rationales can contain testable claims, but they do not provide a measure of probability calibration.

### 4.4 Information ablations have small endpoint effects

Across 200 ablation campaigns, no single removed channel produced a large or universal loss (Figure 5). Removing diversity reduced D2R recall by 0.016 and was neutral or slightly positive elsewhere. Removing memory changed means by -0.004 to +0.004. Removing uncertainty improved D2R by 0.016, was nearly neutral on Mpro and TYK2, and reduced USP7 by 0.014. Shuffling evidence changed the mean by exactly zero on every target.

**Figure 5:**
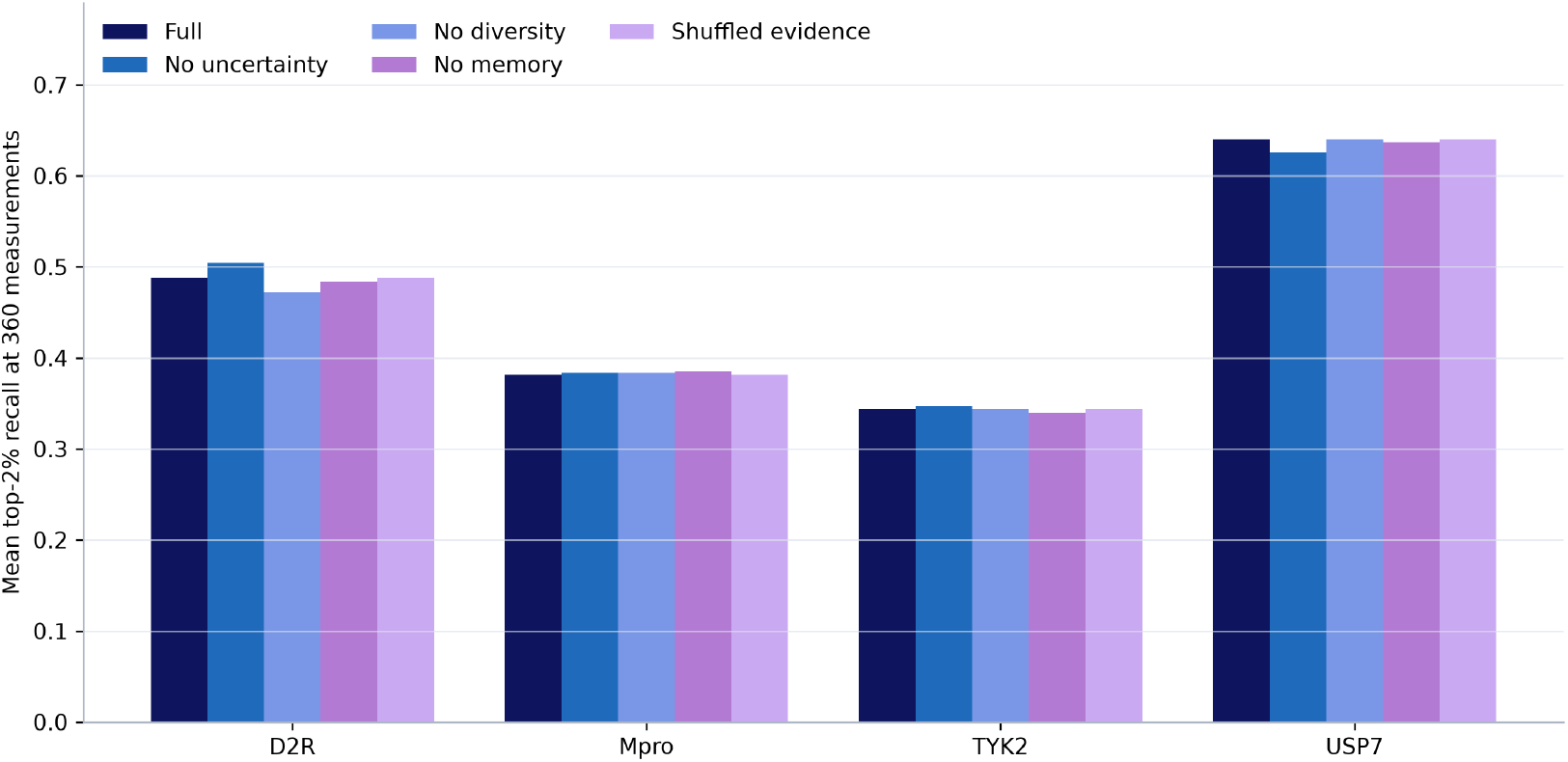
Mean endpoint top-2% recall for the full-information rerun and four paired ablations. Differences are small and target-specific. Full-reference reruns are compared only with their paired ablation arms.

At the decision level, shuffled evidence produced the same allocation path as full information in all D2R, Mpro, and TYK2 campaigns and nine of ten USP7 campaigns. Therefore, fingerprint-bit identities in this evidence packet were usually unnecessary for the observed allocation. The ablation full-reference means differ from the core means because Codex task inference is not a frozen deterministic snapshot; only within-rerun paired comparisons have a causal interpretation.

### 4.5 Availability constraints lower attainable recall

An 80% post-initial availability mask reduced agent top-2% recall by 0.070 on D2R, 0.072 on Mpro, 0.114 on TYK2, and 0.069 on USP7 (Figure 6). Under this constraint, the agent exceeded the strongest constrained fixed policy on Mpro and USP7 but remained behind on D2R and TYK2. The strongest fixed policies also lost recall relative to their unconstrained counterparts, confirming that access changes the problem rather than only disadvantaging the agent. Across 12,000 post-initial agent selections, no unavailable compound was selected.

**Figure 6:**
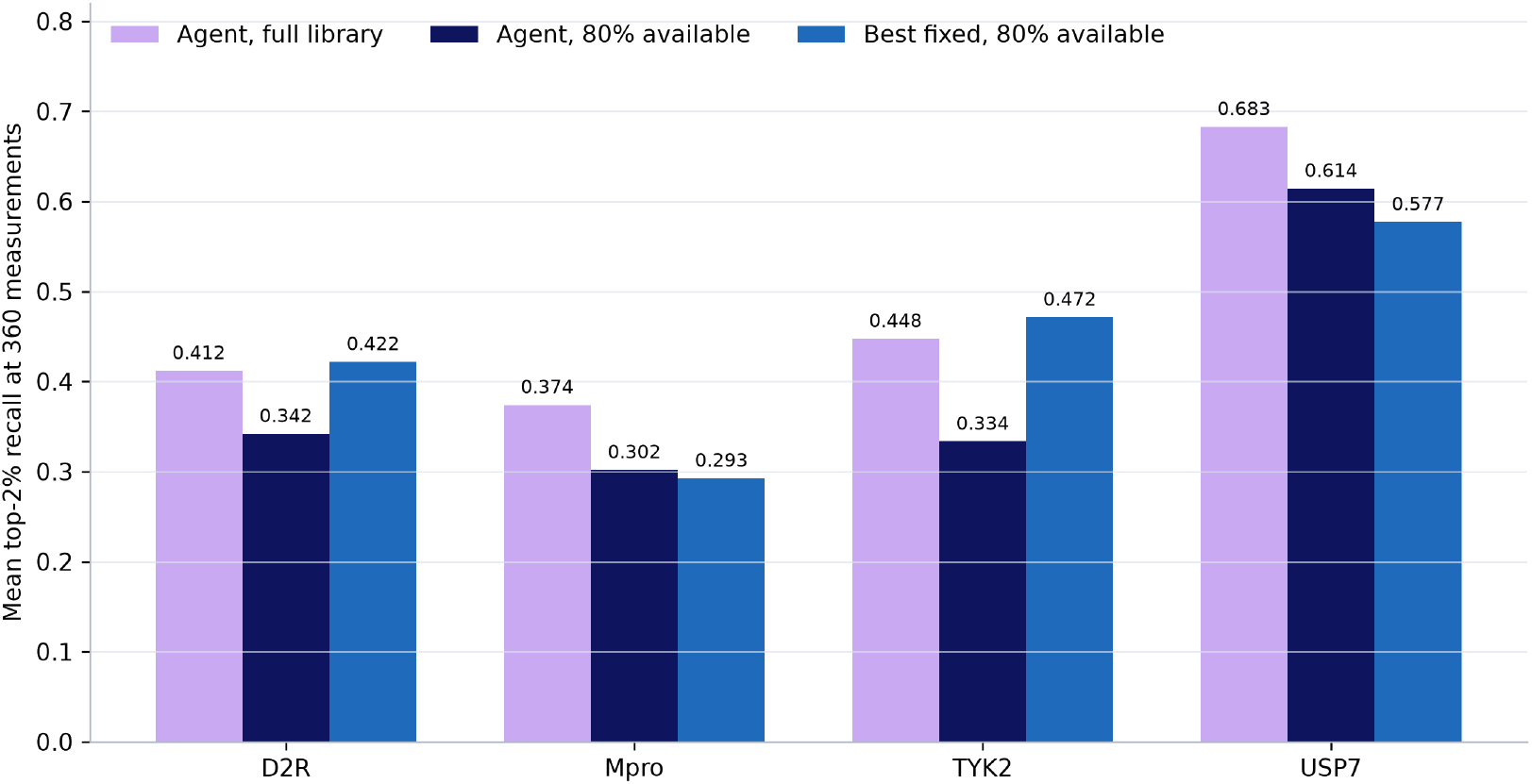
Mean endpoint top-2% recall for the unconstrained agent, the 80%-availability agent, and the strongest constrained fixed policy. Each bar averages ten paired seeds.

### 4.6 Endpoint sensitivity across recall thresholds

The target-level ordering remains visible when the endpoint threshold changes from the top 1% to the top 5% of compounds (Figure 7). USP7 has the highest mean recall at every threshold, whereas Mpro and TYK2 show lower top-5% recovery than top-1% recovery because the broader valuable set is harder to cover within the same assay budget.

**Figure 7:**
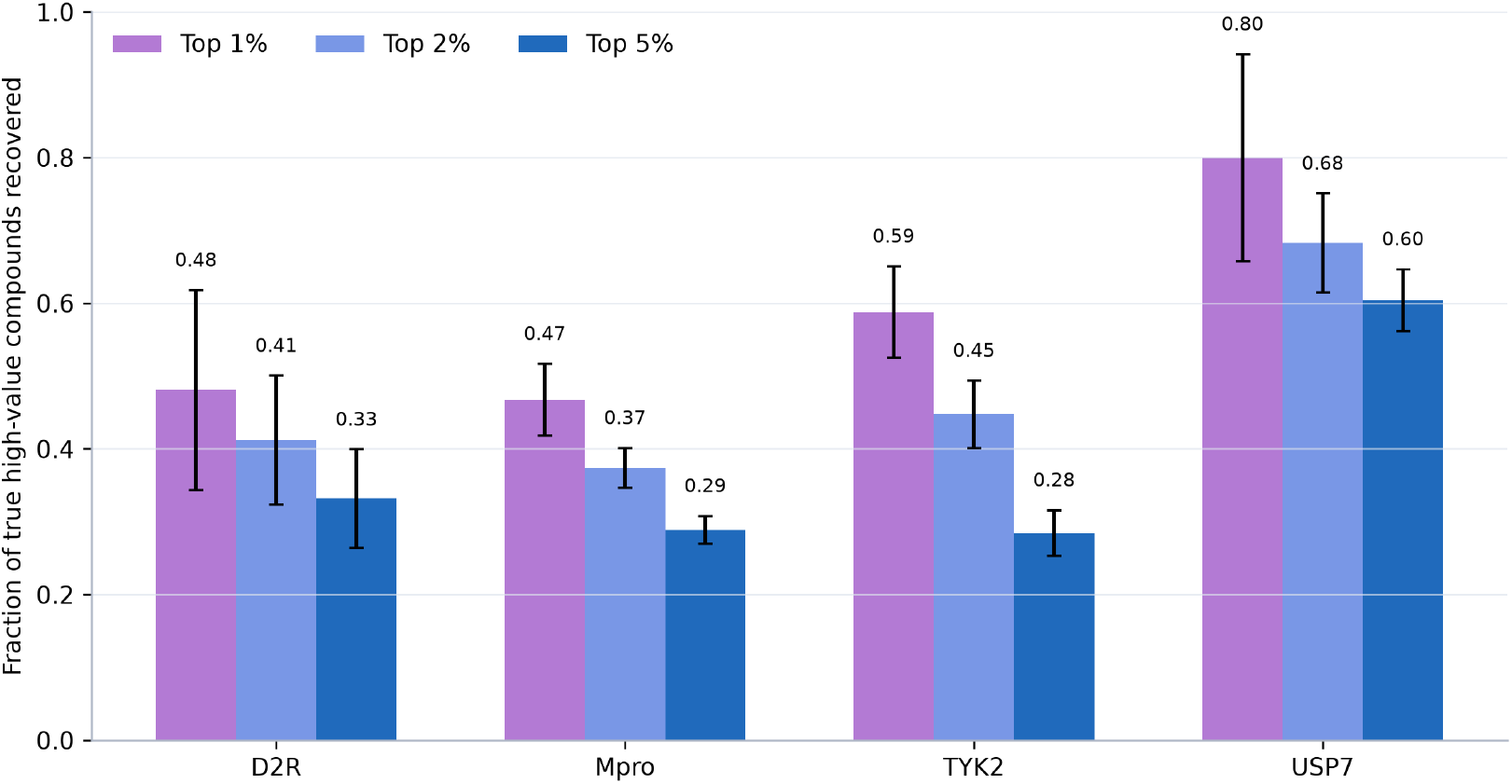
Agent endpoint recall at the top-1%, top-2%, and top-5% thresholds. Error bars show standard deviation over ten seeds.

## 5 Discussion

The benchmark supports three conclusions. First, language-model strategy control is useful: it beat random in every matched campaign, respected every batch constraint, and found the global optimum reliably on three targets. Second, adaptation is not the same as dominance. Target-specific fixed policies were better on D2R and TYK2, and the agent’s allocations reveal why: TYK2 rewarded aggressive exploitation, while D2R favored a stable balanced mixture. Third, explanation channels require causal testing. High threshold-claim success shows that rationales can be empirically checked, but the shuffled-evidence ablation shows that one ostensibly explanatory channel rarely changed actions.

These findings argue for agents that are evaluated as decision policies rather than judged from fluent rationales. A practical agent should state quantitative predictions, expose its allocation, obey feasibility constraints, and be compared with strong target-specific policies. Future versions should emit probabilities for formal calibration and use a frozen versioned API snapshot when exact replication and token accounting are required.

### Limitations

The study is retrospective and uses known affinity labels as a virtual assay; it does not model wet-lab noise, synthesis time, procurement cost, or assay failure. The GP, Tanimoto kernel, and fingerprint representation are fixed for fairness, so conclusions may change with other surrogates. Codex-task inference is not an exact model snapshot. The threshold scorer captures explicit templates but not every qualitative claim. Finally, four targets are enough to expose heterogeneity but not to establish broad generality.

## 6 Conclusion

A language-model agent can successfully allocate exploit, uncertainty, and diversity queues during batch ligand discovery. It consistently improves over random sampling and can match the strongest fixed policy on some targets, but it is not universally superior. Paired ablations and availability constraints show that both information use and operational access must be tested directly. The most defensible role for the language model is therefore an auditable strategy layer inside a numerically controlled active-learning system.

